# Using *Galleria mellonella* to study pathogen dissemination and anti-infective surface coatings

**DOI:** 10.1101/2024.06.11.598455

**Authors:** Evgenia Maslova, Ciaram Staber, Ronan R McCarthy

## Abstract

Hospital associated infections and localised hospital outbreaks are a major challenge for infection control teams (ICTs) in hospitals around the world. Advances in artificial intelligence and infection modelling have enabled ICT teams to better predict and trace infection spread. However, it is notoriously difficult to replicate bacterial dissemination or validate prediction models in a laboratory setting due to a lack of effective *in vivo* models. In this work, we sought to develop *Galleria mellonella* as a model organism to replicate the dissemination of pathogens in a hospital intensive care unit (ICU). By combining this model organism with 3D printed models of a real hospital ICU, we are able to demonstrate that larvae do disseminate the multidrug resistant pathogen *Acinetobacter baumannii* within this ICU model and that it is possible to use this model to identify infection hotspots. Importantly, this model can also be used for intervention strategy testing as we also show that bacterial dissemination can be significantly mitigated by the introduction of antimicrobial wall surface coatings. This model provides a robust platform for the testing of antimicrobial surface coatings as well as the study of genetic determinants with a role in pathogen dissemination.

## Introduction

Controlling the spread of nosocomial pathogens within the hospital-built environment is a major challenge for infection control teams (ICT), particularly as many of these nosocomial pathogens are multidrug resistant. *Acinetobacter baumannii* is a Gram-negative opportunistic pathogen primarily associated with causing nosocomial infections. *A. baumannii* is recognised as the World Health Organisations number one priority pathogen due to its capacity for causing localised hospital outbreaks, its ability to survive on hospital surfaces for prolonged periods and its recalcitrance to antibiotic therapy (Doughty *et al*., 2023, Green *et al*., 2022, J.D. Hunter, 2012, Ibrahim *et al*., 2017, Fernández-Billón *et al*.,2023, Pachori, Gothalwal,&Gandhi’s, 2019, Chen *et al*. 2022).

Developments in sampling, sequencing and tracking technologies have made it possible to visualise and analyse pathogen spread in hospital settings in near real-time (Haverkate *et al*., 2015, Nguen et al., 2020, King *et al*., 2021, Doughty *et al*., 2023). Modelling is also gaining an increasing importance in uncovering how these pathogens disseminate. Several *in situ* simulations have been developed in order to visualise and predict pathogen dissemination in hospital wards (Gibbs *et al*., 2018, Kotay *et al*., 2017). However, the majority of simulation models such as agent-based model (ABM), system dynamics model (SD) and hybrid simulation models do not facilitate prevention strategy testing nor do they typically facilitate the study of different strains of the same species (Nguen *et al*., 2020). The further experimental validation of these simulation models is often not feasible as there is a lack of affordable *in vivo* dissemination models available.

In this study, we propose a bacterial dissemination model utilising *Galleria mellonella* as the surrogate vector for pathogen spread. *G. mellonella* is a robust model organism that has gained popularity in recent decades. This model organism has been used for a wide range of assays including virulence assays (bacterial and fungal), immunotoxicity testing (concerning the adaptive immune system), drug toxicity assays, testing the effects of ultraviolet radiation, probiotics, and bacteriophage treatment efficacy (Cunha *et al*., 2023, Stempinski, Smith and Casadevall 2022, Erbas *et al*., 2023, Demirtürk, Uçkan, and Mert, 2023, Sabockyte *et al*., 2023, Filippov *et al*., 2022, Maslova *et al*., 2023). *G. mellonella* are affordable, have low ethical considerations associated with their use and have low maintenance and handling requirements which could facilitate faster early-stage pathogen dissemination and prevention strategy testing. Here, we demonstrate the capacity of a *G. mellonella* larvae-based model to be used as a tool to study bacterial dissemination and to evaluate the efficacy of antimicrobial surface coatings *in vivo*.

## Results

### *G. mellonella* act as vectors to disseminate bacteria throughout a 3D printed ICU model

Hospitals are potent reservoirs for several opportunistic multidrug resistant bacterial pathogens. Due to the nature of the healthcare environment, pathogens can disseminate via a multitude of routes and agents within the hospital setting, with humans being one of the primary vectors for infection spread (Webb *et al*., 2005, Hassoun-Kheir *et al*., 2020). Here, we proposed an ICU ward model utilising the *G. mellonella* as a surrogate for humans as vectors of pathogen dissemination. To add additional clinical relevance to our model, we designed a replica hospital ward based off publicly available schematics of Brigham and Women’s Hospital ICU (MA). The model included 11 patient rooms, a hallway and a central nurses station (Figure 1A). The 3D printed model was sterilized using an IMS solution and tested to confirm that the larvae can move freely around the model ICU (Video 1). We then tested whether the larvae could spread bacteria around the hospital by placing the model ICU on a bed of LB agar and subsequently introducing a single site of bacterial colonisation in Room 13 which is the central nurses station. This location was chosen due to the previous reports highlighting non-treatment areas such as doctor and nurse stations as important environmental reservoirs for bacteria (Dumford *et al*., 2009, Bures *et al*., 2000, Tajeddin *et al*., 2016). We chose to inoculate the ICU with a sample of *A. baumannii* AB5075. *A. baumannii* is a highly prevalent nosocomial pathogen commonly associated with ICU outbreaks around the world (Wieland *et al*., 2018). A single larva was then added to the model and allowed to move freely through the bays of the model for 16-18 hours. Based on visual observation of the agar bed underneath the ICU model, it was clear that bacteria had been spread throughout every room in the ICU (Figure 1C).

**Figure 1.**
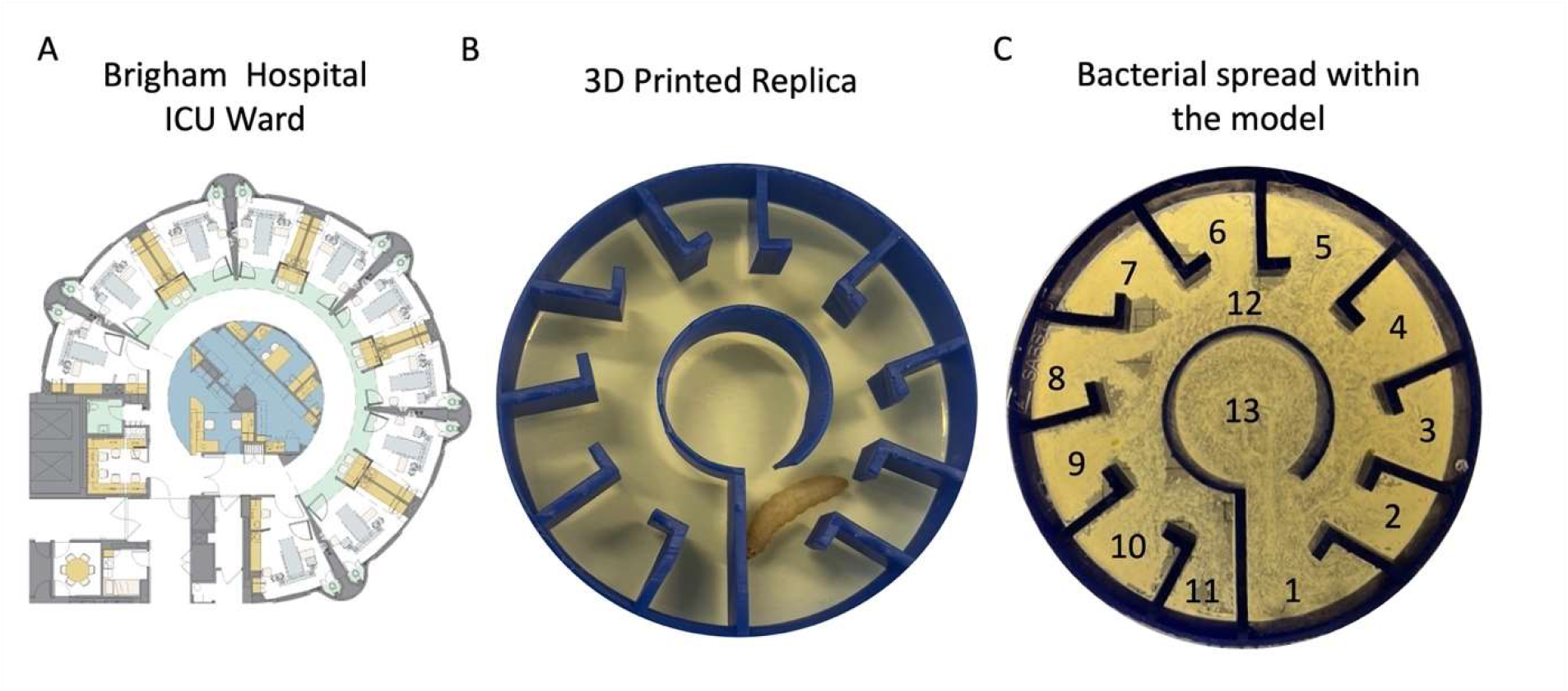
The experimental design and layout of the hospital ICU ward. (A) The outline of the hospital ICU model. (B) The experimental set-up with the 3D printed ICU model placed upon a bed of LB agar. A piece of agar seeded with *A. baumannii* AB5075 can be seen in Room 13 (Central Nurses Station). (C) The bacterial dissemination by a singular *G. mellonella* larva can be seen as growth on an LB agar after 16-18h incubation *A. baumannii* AB5075.

To develop the model further, we replaced the LB agar bed (which facilitates bacterial growth) with Whatmann paper and included 11 larvae in the model (one for each of the ICU rooms with the exception of the nurses station and the hall) (Figure 2A). To enable the quantification of bacterial dissemination, after a 24-hour incubation period, the walls of each room were swabbed and the colony forming units (CFUs) from each enumerated (Figure 2B). The nurses station (Room 13) reproducibly returned the highest CFU values as expected, as it was where the initial starting inoculum was located. Rooms 1-5, 7 and 12 had higher CFU levels than the rest of the rooms, which was expected as Room 12 is the “hallway” of the ICU model and would therefore have the highest visit count. Rooms 1-5 are the first rooms the larva would encounter once exiting Room 13 which also explains the high CFU counts in these rooms. Rooms 6-7 and 8-11 had the lowest number of bacteria recovered. This data demonstrates that the model can be used to study bacterial dissemination throughout a replica ICU and to identify potential hotspots for bacterial contamination.

**Figure 2.**
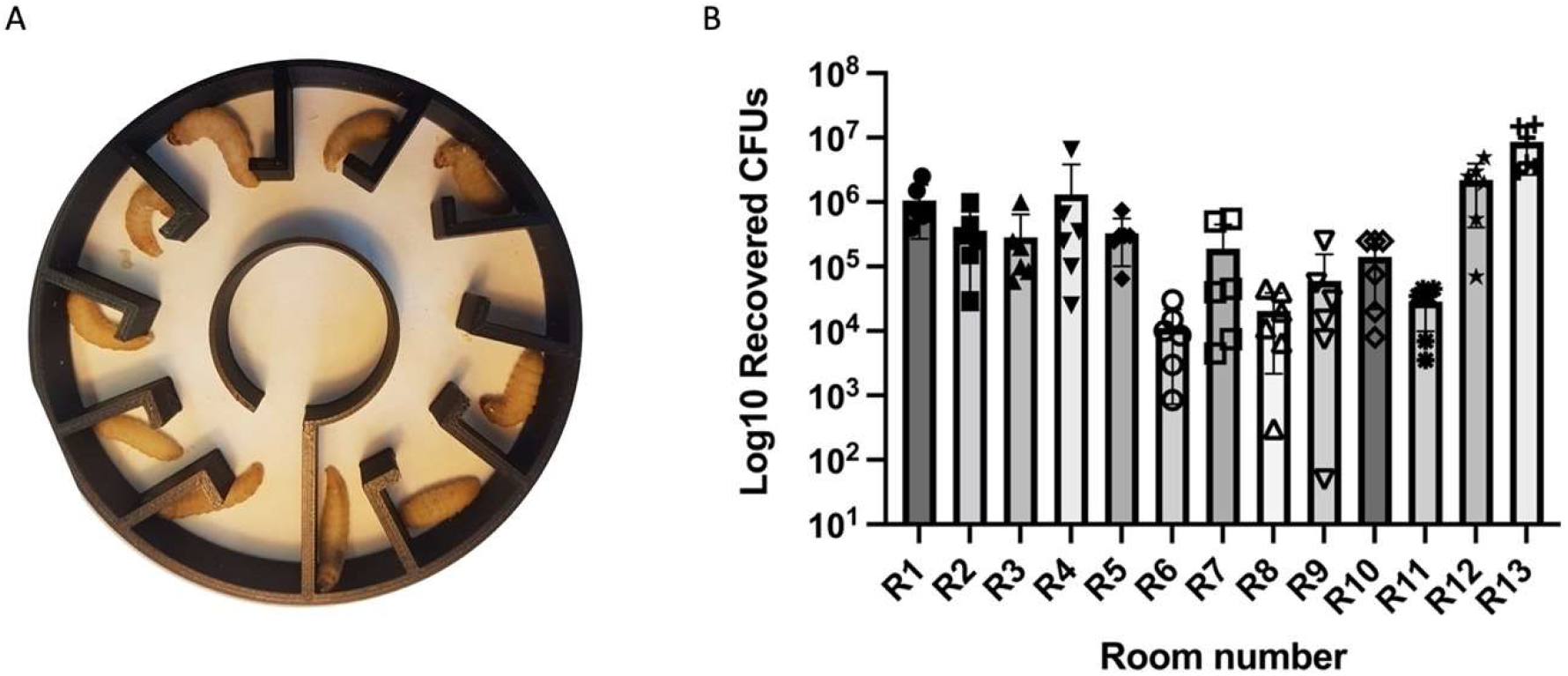
Uncoated ICU model set up with 11 larvae (A) and the average *A. baumannii* AB5075 CFU values obtained from each room. **(B)**. This data represents the average CFUs obtained from 6 biological replicates after 24h incubation with average shown with SD. Ordinary one-way ANOVA with Tukey correction was performed, p value < 0.0001.

### Surface coating decreases bacterial bioburden on the model surfaces

Once the bacterial load distribution within the ICU model was confirmed, we next sought to explore the capacity of the model to be used to evaluate antimicrobial surface coatings. Antimicrobial surface coatings are commonly used to limit microbial surface contamination and represent a potential strategy to limit the spread of pathogens within a hospital. We developed a PVA based paint solution supplemented with gentamicin as the antimicrobial agent (Menetrez *et al*., 2008, Fulmer *et al*,, 2011, Ren *et al*., 2022). Gentamicin was chosen for its ability to maintain potency even after being exposed to the high temperatures needed to dissolve the polymers. In the model with the gentamicin loaded surface coating, less bacteria were recovered in every single room as compared to the models with no coating, with statistically significant reductions seen in rooms 1, 2, 5, 8, 10, 12 and 13 (Figure 3,4). Overall, this demonstrates the capacity of this model to effectively determine the efficacy of antimicrobial surface coatings *in vivo*.

**Figure 3.**
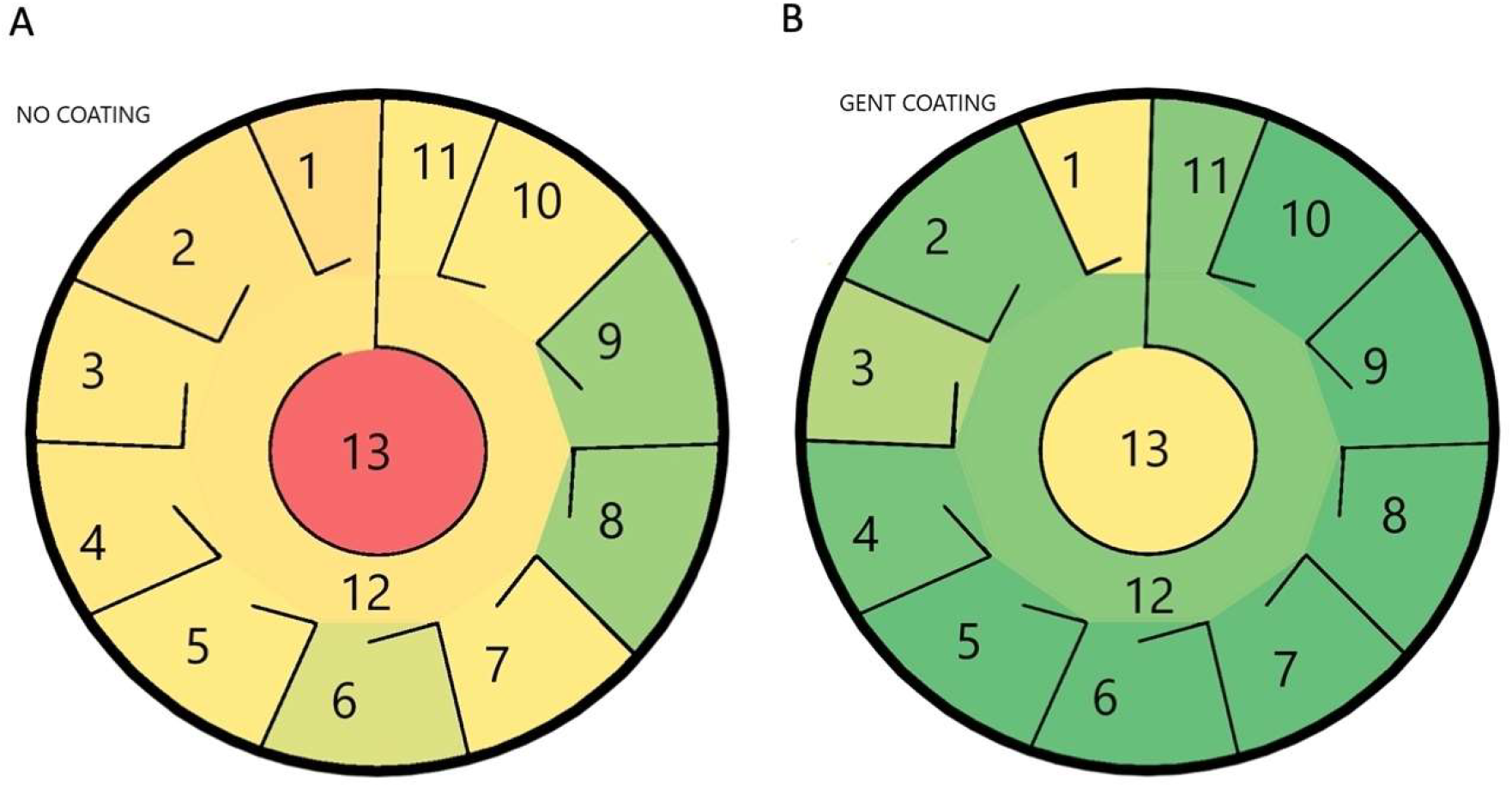
The bacterial spread by *G. mellonella* in the non-coated and gentamicin coated ICU models. The bacterial spread of *A. baumannii* with uncoated (A) and gentamicin (B) coated model respectively. The color coding corresponds to the highest (red), the medium high (yellow) and the low (green) bacterial densities. Room 13 was the infection start site and has the highest CFU in both ICUs.

**Figure 4.**
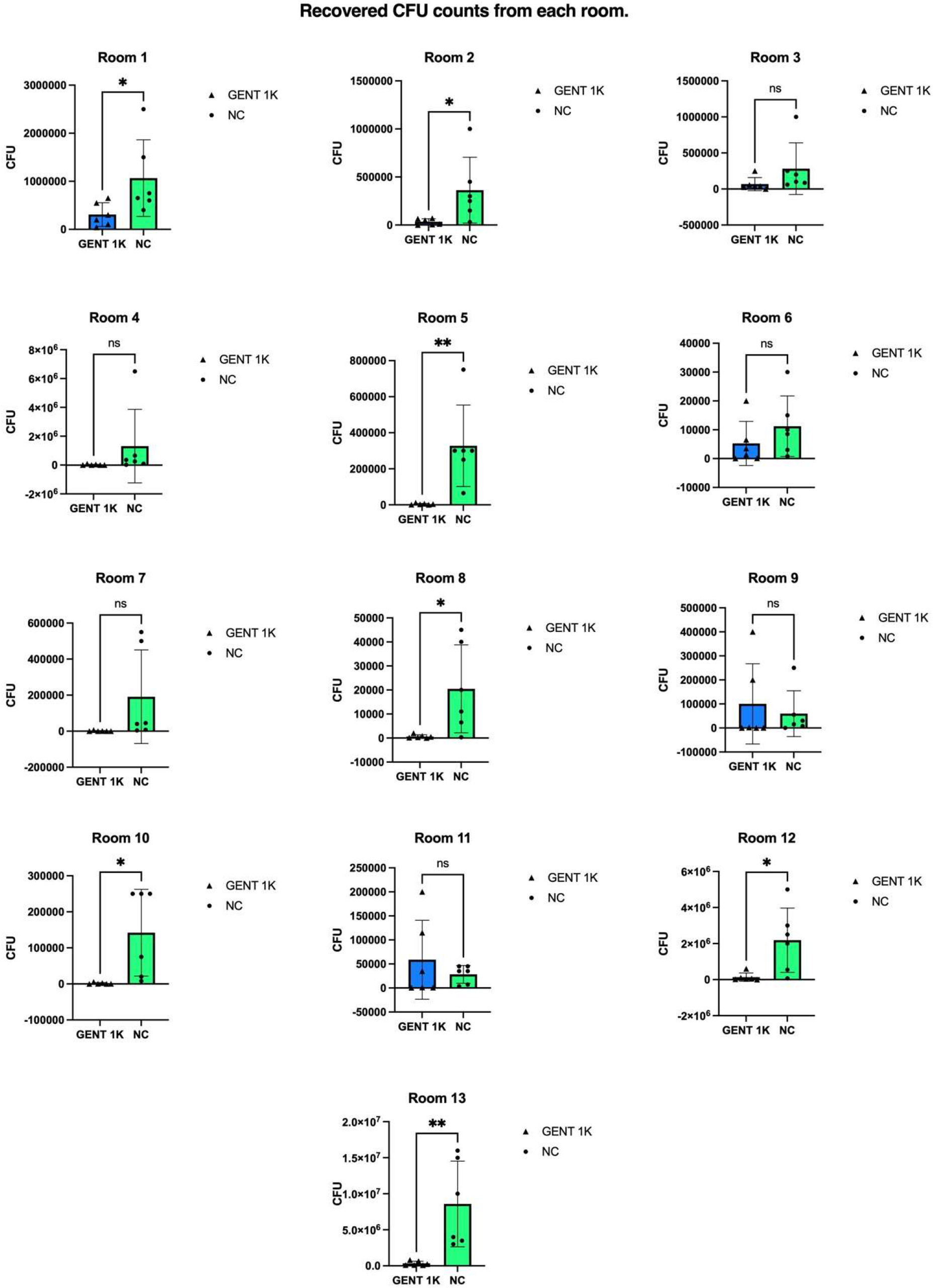
The CFU counts recovered from each room of the gentamicin coated and non-coated ICUs models. Gentamicin treated rooms exhibited a consistent reduction in CFUs with the exception of rooms 9 and 11. Rooms 3,4,6,7,9,11 exhibited no significant decrease in CFUs. This experiment has 6 biological repeats. Unpaired student t-test not significant p value was >0.05, * p value <0.05, ** p value < 0.005.

## Discussion

Amidst the multidrug resistance crisis, understanding the dissemination of nosocomial pathogens through the hospital-built environment is crucial. Here, we developed a bacterial dissemination model utilizing 3D printing and *G. mellonella* as the pathogen dissemination vector. This model was tested with *A. baumannii*, a notorious nosocomial bacterial pathogen with a remarkable capacity to survive on surfaces for prolonged periods. It was shown that the larvae could traverse the entire hospital ICU ward model within a 16-18h window, spreading bacteria to the entire ICU in the process (Figure 1C). This can be viewed as a pathogen starting in one bay then possibly being spread to others via an individual commuting between bays in the ward while not adhering to appropriate safety and infection control protocols. Having shown the proof-of-principle that one larva was able to spread bacteria throughout the ICU, the bacterial spread was quantified in a multi-vector experiment. This would allow us to determine what areas were at a higher risk. The amount of larva used per Petri dish was increased to match the number of outer rooms being 11. As expected, the highest number of CFUs were recovered from the center room which is labelled Room 13 (Nurses station) (Figure 2B). Rooms closest to the entrance of Room 13 typically had higher CFU numbers. These being Rooms 1-5 and 12 with 12 being the central corridor connecting all rooms. Rooms that were further from that entrance like 9, 10 and 11 showed lower CFUs. This trend shows that the CFU recovered generally goes down from room to room with increasing distance from the start site. While in this study we used insects as surrogates for human pathogen spread, several studies have also demonstrated the relevance of insects such as cockroaches and ants as vectors/reservoirs for multidrug resistant bacteria in the hospital environment highlighting their role in nosocomial infection spread (Fowler *et al*., 1993, Do Nascimento *et al*., 2020, Abdolmaleki *et al*., 2020, Menasria *et al*., 2014, Hassan *et al*., 2021).

With confirmation of the bacterial spread by *G. mellonella* throughout the model, further testing was done to determine if this model could be used to test potential antimicrobial agents. The PVA base polymer coating was chosen due to its low costs and low toxicity (DeMerlis and Schomeker, 2003). PVA is already well established within the pharmaceutical and medical industry for a wide array of applications. Across all experiments the results showed that a gentamicin coating was able to reduce the CFUs recovered (Figure 3,4). Clearly, the use of antibiotics directly within a surface coating in everyday use is not realistic as it would contribute to further pressures for antibiotic resistance. However, as a proof of principle it validates the capacity of this model to be used as an *in vivo* tool for testing the efficacy of antimicrobial surface coatings. Such a model has the potential to accelerate the development of these technologies and add further relevance to *in silico* models that already exist in the field.

The experimental set-up is highly versatile, for example bespoke 3D models can be designed to give representative insights for a specific hospital, the initial inoculation site can be altered, the number of larvae/ vectors can be modified. Equally the options for surface coatings can be modified to address a specific research question, testing for example different doses or formulations. Equally, different pathogens, polymicrobial communities, surface biofilms can all be tested in the model. The is also the possibility that the surface of the model can be washed and the assay can be repeated to determine the durability and leeching of any surface coating. The model can also be run at scale, facilitating the generation of large volumes of data that can be used to train artificial intelligence applications or to test *in silico* dissemination models. A key consideration however is that the movement of the larvae is more random and less directed as compared to the movements of hospital staff on a ward, however the same limitation is likely to apply to any animal used in a hospital dissemination model study.

The model also has the potential to be used to study the genetic determinants of infection dissemination as different strains can be used to inoculate the model and their relative capacity for dissemination and persistence can be determined. Equally, if specific genes have been hypothesized as being linked to transmissibility or built environment persistence, it will be possible to use this model to gain *in vivo* insights about the consequences of mutating or overexpressing these genes. In conclusion, this study demonstrates *G. mellonella* ICU model can be used to study pathogen dissemination and the capacity of surface coatings to limit this dissemination.

## Materials and methods

### Bacteria strains and growing conditions

*A. baumannii* AB5075 was grown overnight at 37°C in 5ml of LB broth at 180 rpm for 16-18h. To obtain the lawn, 100 µl of the AB5075 overnight culture was put onto a nonselective LB agar plate and spread evenly across the surface using a sterile swab or inoculation loop. Then grown at 37°C for 16-18h. Incubation of the single larva experiment set up with the patch of bacteria lawn was for 16-18h at 37°C, and 24h for the set up with 11 larvae. Once samples were collected, serially diluted, and plated they were incubated in the static incubator for 16-18 h then counts were taken.

### Animal acquisition and preparation

*G. mellonella* were sourced from LiveFood UK Ltd. (Somerset, United Kingdom). The larvae were stored at 4°C in wood shavings upon arrival until use. During the selection process, the larvae were checked to ensure there was no signs of melanization along its body as it is a sign of an infection or trauma in *G. mellonella* (Pereira *et al*., 2018). Once selected, the larvae were sterilized by rolling the larva in a shallow pool of 70% IMS in a sterile Petri dish. The larva was then dried off by putting it on a sterile piece of blue paper towel roll. The larvae were then moved to a sterile Petri dish and incubated at 4°C until needed. Due to the storage at 4°C, the larvae remain immobile until they warm up to the room temperature which facilitates easier sterilization and handling of the larvae during the experiment set up.

### 3D printed ICU model

The schematic ICU model was made using a resin (PVA) and a 3D printer (Elegoo Mars 2 Pro). It was modelled off of an intensive care ward of Brigham and Womans Hospital, MA, USA. The model was stored in a dry cool place when not in use and was sterilised by soaking it in 70% industrial methylated spirits (IMS) for a minimum of 90 seconds prior to use. It was then moved to sterile Petri dish and allowed to air dry in a laminar flow hood. Once dry, it was placed on top of the LB agar media for the single larva experimental set up. For the 11 larvae set up, no agar was introduced into the Petri dish and layers of foil were used to elevate the ICU until it met the lid of the Petri dish when closed, preventing larvae from climbing over the walls. Whatmann filter paper was then used to separate the foil layers and the ICU model.

### ICU model bacterial dissemination assay set-up with a single larva

10 ml of lysogeny (LB) agar was poured into a sterile Petri dish. Once dried, the sterile ICU model was placed on top of the agar. The Petri dish was then covered, and the lid gently pressed to implant the ICU model onto the agar. A 50µl droplet of *A. baumannii* AB5075 overnight culture adjusted to OD_600_ = 1 was then pipetted into the central Nurses Station referred to as Room 13. A single larva was then placed into same room as the bacteria. The set-up was incubated at 37°C for 16-18 h. After the incubation period, the larva was euthanized by placing it at -20°C. The ICU model was removed from the agar and the bacterial spread left by the larva in the Petri dish was assessed.

### ICU model bacterial dissemination set up used to obtain bacterial load data from the walls of the ICU

A 1 cm^2^ patch was cut out of an LB agar plate that was seeded with a lawn of *A. baumannii* AB5075 on it. The patch was then placed into the Nurses station of the ICU replica. Sterilised larvae were then added to each outer room of the ICU. Tape was used to secure the lid to the plate so that there was no gap between the walls of the ICU effectively making the lid of the Petri dish, the ceilings of the ward. The Petri dish was then be incubated for 24h at 37ºC.

### ICU model walls CFU sample collection and evaluation

At 24 hours all larvae were moved to -20°C and euthanized. The ICU model was then moved to a new sterile Petri dish using sterile tweezers. A sterile PBS-soaked swab was then used to sample each room, moving to swab across all walls. The swab’s tip was broken off into an Eppendorf tube with 500 µl PBS and vortexed. This was repeated for each room including the Nurses Stations (central room, Room 13) and the corridor. The samples were then serially diluted to 10^-5^ in PBS and plated onto LB agar supplemented with 50 µg/ml kanamycin, to limit any background growth. The plates were incubated for 16-18h. Post incubation, individual colonies were counted and used to calculate the colony forming units (CFUs) of each room.

### Surface coating mixture composition and preparation

The surface coating solution was made using the composition proposed by Marcelo *et al*. (2022). The polyvinyl alcohol (PVA):polyvinylpyrrolidone(PVP) polymer solution was made by dissolving 2000 mg of PVA and 500 mg of PVP in 15 mL of distilled water. The mixture was then incubated in the 80°C water bath, occasionally being vortexed until the solution went from cloudy to transparent. Using this solution as a base coating solution, the total volume would be used to make % w/v treatment solution mixtures. Gentamicin was chosen for its ability to maintain potency even after enduring the high temperature needed to dissolve the polymers. Gentamicin 16 mg/ml was used as the positive control as this above the EUCAST breakpoint for *A. baumannii*.

### Coating the ICU surface

After undergoing sterilisation and drying the ICU model would be coated with the desired mixtures containing the active ingredient. This was done by saturating a cotton swab in 1 ml of the solution. The swab was then used to coat the outer walls of each room 1-12. The same swab was remoistened and used to coat the inner and outer wall of the central room. This was repeated until virtually all the 1 ml of solution was used up whilst re-saturating the swab every 3-4 rooms. Only one swab was used to coat each ICU model. The same procedure would then be followed as described in the infection spread set up section mentioned above, but with the coated ICUs.

### Statistical analysis

For analysis of the total CFU recovery of the ICU models the one-way ANOVA with Tukey correction test was used. Analysis of the antibiotic coating solution efficacy to reduce bacterial proliferation was done using unpaired student t-test.

## Data Availability

All data including the 3D printed model of the ICU are available from the corresponding author.

